# How neurons move during action potentials

**DOI:** 10.1101/765768

**Authors:** Tong Ling, Kevin C. Boyle, Valentina Zuckerman, Thomas Flores, Charu Ramakrishnan, Karl Deisseroth, Daniel Palanker

## Abstract

Neurons undergo nanometer-scale deformations during action potentials, and the underlying mechanism has been actively debated for decades. Previous observations were limited to a single spot or the cell boundary, while movement across the entire neuron during the action potential remained unclear.

We report full-field imaging of cellular deformations accompanying the action potential in mammalian neuron somas (−1.8nm~1.3nm) and neurites (−0.7nm~0.9nm), using fast quantitative phase imaging with a temporal resolution of 0.1ms and an optical pathlength sensitivity of <4pm per pixel. Spike-triggered average, synchronized to electrical recording, demonstrates that the time course of the optical phase changes matches the dynamics of the electrical signal, with the optical signal revealing the intracellular potential rather than its time derivative detected via extracellular electrodes. Using 3D cellular morphology extracted via confocal microscopy, we demonstrate that the voltage-dependent changes in the membrane tension induced by ionic repulsion can explain the magnitude, time course and spatial features of the phase imaging. Our full-field observations of the spike-induced deformations in mammalian neurons opens the door to non-invasive label-free imaging of neural signaling.

## Introduction

Minute (nanometer-scale) cellular deformations accompanying the action potential have been observed in crustacean nerves (0.3~10 nm)^1–4^, squid giant axon (~1 nm)^5,6^ and, recently, in mammalian neurons (0.2~0.4 nm)^7,8^. These findings illustrate that rapid changes in transmembrane voltage due to the opening and closing of the voltage-gated ion channels have mechanical consequences accompanying action potentials and other changes in cell potential, which may allow non-invasive label-free imaging of neural signals^9^. However, detecting the nanometer-scale millisecond-long deformations in biological cells is very challenging, and previous observations were limited to a single spot on the axon^1–5,7^ or to the boundary of the cell soma^8^. Until now, the entire picture of how neurons move during the action potential was unclear, and the underlying mechanism remains poorly understood. Several hypotheses have been proposed to explain the membrane displacement during the action potential in axons, including electrostriction^10^, converse flexoelectricity^11,12^ or soliton wave^13,14^. Each of these mechanisms were shown to fit the time-course of the reported mechanical displacements, precluding a definitive answer to what the actual mechanism is.

Not only neurons, but also other cells, such as HEK-293, deform when their membrane potential is altered^9,15–19^. Atomic force microscopy showed that the membrane displacement in HEK-293 cells is proportional to the membrane potential, and a hypothesis was put forth that such displacements are due to voltage-dependent membrane tension originating in the lateral repulsion of mobile ions^15^. This linear dependence was later confirmed by measurements using piezoelectric nanoribbons^16^, quantitative phase imaging (QPI)^18^ and plasmonic imaging^19^. Under this hypothesis, changes in the membrane tension with varying cell potential lead to a pressure imbalance on the cell surface, as described by the Young-Laplace law, thus deforming the cell into a new shape balancing the membrane tension, hydrostatic pressure and the elastic force from the cytoskeleton. Careful analysis of this proposed model calls for a full-field imaging of the whole cell with sub-millisecond temporal resolution and angstrom-scale membrane displacement sensitivity.

In this study, we report the full-field imaging of neuron deformations accompanying the action potential (termed ‘spike-induced deformation’) using ultrafast quantitative phase imaging (QPI)^9,20^ with spike-triggered averaging (STA), which enables 0.1 ms temporal resolution and an axial pathlength sensitivity of 4 pm/pixel. Synchronization was performed first via multielectrode array (MEA) detecting spontaneous firing in real time, and subsequently, for increased numbers of observations, by synchronizing the QPI to light pulses in optogenetic stimulation^21,22^. To evaluate the membrane-tension based model as the underlying mechanism, we compare the experimental results to a proposed model of spike-induced deformation by applying the voltage-dependent membrane tension to the ‘cortical-shell liquid-core’ mechanical model^23–25^, using the 3D cellular morphology mapped by confocal imaging.

## Results

### Noise reduction via spike-triggered averaging (STA)

In addition to offering intrinsic phase contrast of biological cells without exogenous markers, QPI provides an exquisitely precise quantitative metric for the optical path difference^26^, which allows full-field imaging of the nanometer-scale membrane displacements along the optical axis. In our system, a transparent MEA with cultured neurons was mounted on the QPI microscope and illuminated by a collimated beam incident from the top. Cellular deformations change the incident wavefront distortions, resulting in spatial variations of the interferogram imaged by the fast camera (Fig. 1a).

Sensitivity to such wavefront distortions, or optical phase changes, is limited by the shot noise of the detectors, in the absence of other more dominant interference^27^. Given the 11,000 electron well capacity in our camera, the phase noise from a single shot in our system is ~2 mrad, which is ~60 times larger than the sensitivity needed to image the nanometer-scale spike-induced deformation in transmission geometry^9^. To reduce the phase noise, we implemented spike-triggered averaging (STA). Timing of the electrical spikes recorded by the MEA were used to synchronize the beginning of several thousand phase movies, each recording the full-field phase changes during a single action potential. These movies were synchronized and averaged into a single STA movie (Fig. 1b), where the sub-nanometer spike-induced deformation could be revealed with a temporal resolution of 0.1 ms in a field of view (FOV) of 106 μm × 80 μm (Fig. 1c, Supplementary Movie 1). Assuming a refractive index difference of Δ*n* between the cytoplasm and the extracellular medium, the membrane displacement Δ*h* can be obtained by

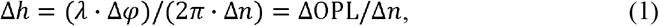

where *λ*, is the center wavelength of the QPI system (819 nm) and ΔOPL is the change in the optical path length.

**Fig. 1.**
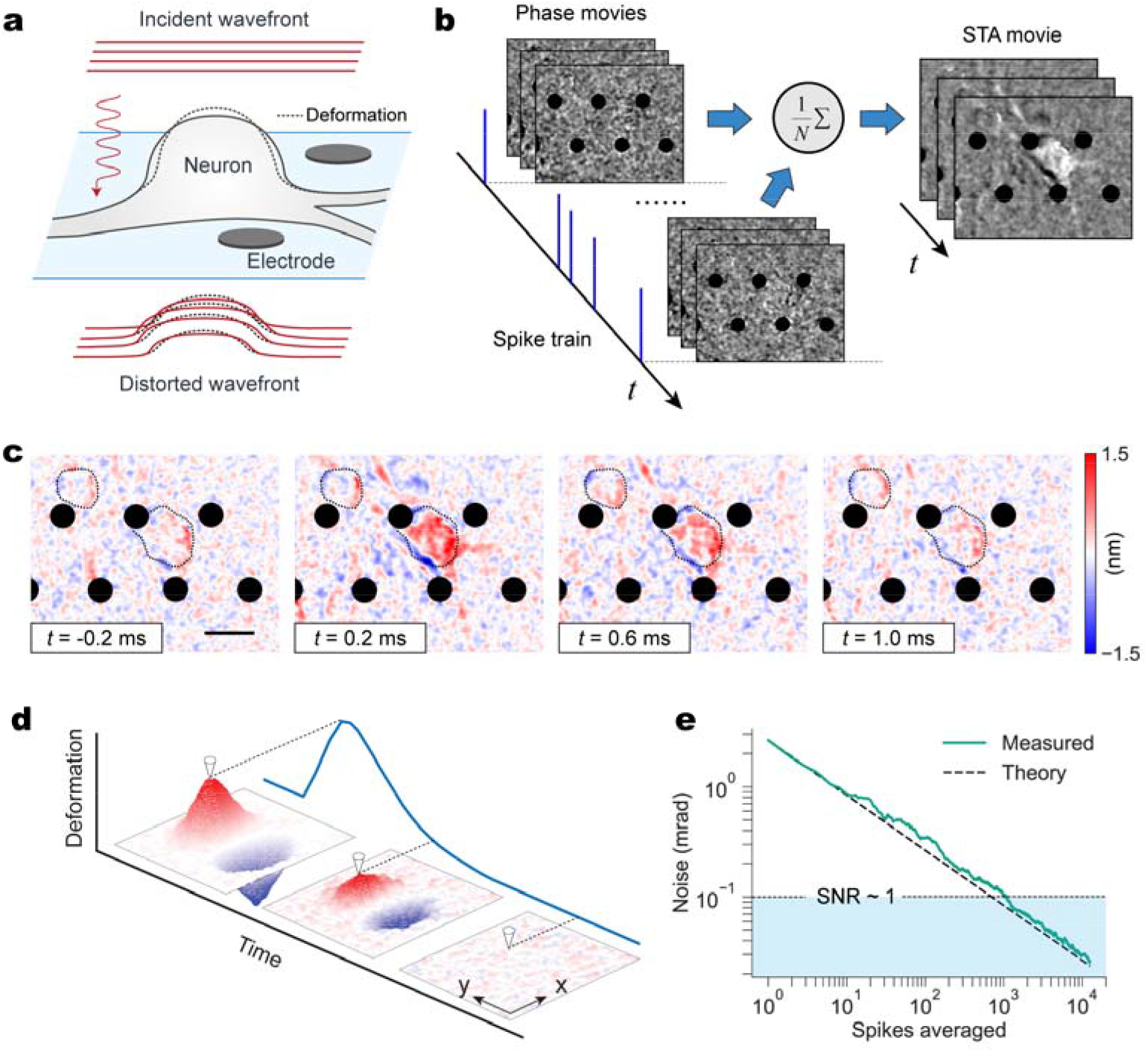
Quantitative phase microscopy combined with microelectrode array recording for spike-triggered average imaging of cellular deformations during action potentials. (a) Collimated incident wavefront accumulates spatially varying phase changes as it passes through cells. Transparent MEA provides simultaneous recording of electrical signals. (b) Spike-triggered average movie is generated by synchronizing the image frames to a preceding electrical spike and averaging multiple events, and electrodes in the movies are masked out with the black dots. (c) Cellular deformations are visualized as the difference between each frame and the initial frame, revealing the nanometer-scale deformation across a neuron. Scalebar: 20 μm. (d) The 4D STAs encode the z displacements of the objects across the field of view, and one point sliced through the marker in time shows the temporal dynamics. (e) Noise in the resulting STAs scales with the number of averages, *N*, according to the theoretical 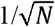.

Ideally, if no correlated noise is present in the STA, the phase noise should scale with the number of averages *N* as 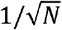, meaning that reducing the phase noise by >60 times requires at least 3,600 averages. Due to the slow, camera-limited data transfer of the very large volume of interferograms (0.5 TB), continuous recording of spontaneous firing yields only 185 seconds of measurements for each hour of operation. At an observed spiking rate of 2~8 Hz, this would yield only 370~1,480 spikes, with 99.2% of the captured data being empty frames without spikes. Therefore, for spontaneously spiking neurons, a crucial step was to detect bursts of activity on the MEA in real time in order to selectively record only the highest density movies to prioritize the limited data storage bandwidth (Supplementary Fig. S1), which resulted in a 4~11 fold increase in the number of averages. With optogenetically controlled recordings, the data collection efficiency improved 10~15 fold since we only captured the 6~10 ms movie segments around each spike time.

The STA movie is a rich 4D dataset that represents the spatial and temporal dynamics of the cellular deformations. Each frame (*t*) encodes deformation (Δ*z*) as a function of position (*x,y*). This is illustrated conceptually in Fig. 1d, and the markers show that slicing through one point in time yields the local temporal dynamics as a plot which can be compared to electrical waveforms recorded via extracellular or intracellular electrodes^9^.

To achieve sub-nanometer sensitivity in the presence of large and ubiquitous interference, including environmental vibrations and particles drifting through the FOV, sophisticated experimental procedures, data filtering and image processing are required. Various corrections, co-optimized between the physical optics and the offline data analysis, resulted in a shot-noise limited performance, which was confirmed by the fact that the noise level scaled with the theoretical 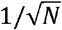 improvement from averaging the uncorrelated noise (Fig. 1e). An optical path length sensitivity of <4 pm per pixel was achieved by the STA with 9091 spikes averaged, corresponding to a membrane displacement sensitivity of <0.2 nm per pixel, given that the refractive index difference Δn is ~0.02 (see Supplementary Materials). Synchronization and phase stability required for averaging without compromising this shot-noise limited performance goes hand-in-hand with the rest of the system design, as described below.

### Spike-induced cellular deformations

We first investigated spike-induced deformations during spontaneous action potentials, in the absence of external stimulation. Primary cortical neurons were cultured on MEAs^28,29^ with a cell density of 1,200~2,000 cells/mm^2^. Starting from ~7 days in vitro (DIV), sparse spontaneous spiking could be observed in electrical recordings. At 17~21 DIV, we recorded the QPI phase movies and electrical signals with real-time burst detection for ~1 hour. In a representative experiment, after averaging over 7,586 spikes, the STA phase movie revealed fast nanometer-scale cellular deformations over the entire soma and neurites during the action potential. Most of the soma rose up, while the left boundary fell as the membrane depolarized (Fig. 1c, Supplementary Movie 1).

Spatial averaging of the phase signal over the soma, considering the polarity of the phase shift in each area^9^, improved the peak signal-to-noise ratio (SNR) to 21.5, yielding a high-fidelity time-course of the phase changes including a quick onset (~0.2 ms) to 0.12 mrad followed by a ~1 ms relaxation (Fig. 2a). The time derivative of the phase signal closely matches the electrical waveform on extracellular electrodes, which corresponds to the time derivative of the membrane potential with an inverted sign (Fig. 2a). Latency between the phase signal and electrical signal was not detected, which means it was below our time resolution of 0.1 ms.

**Fig. 2.**
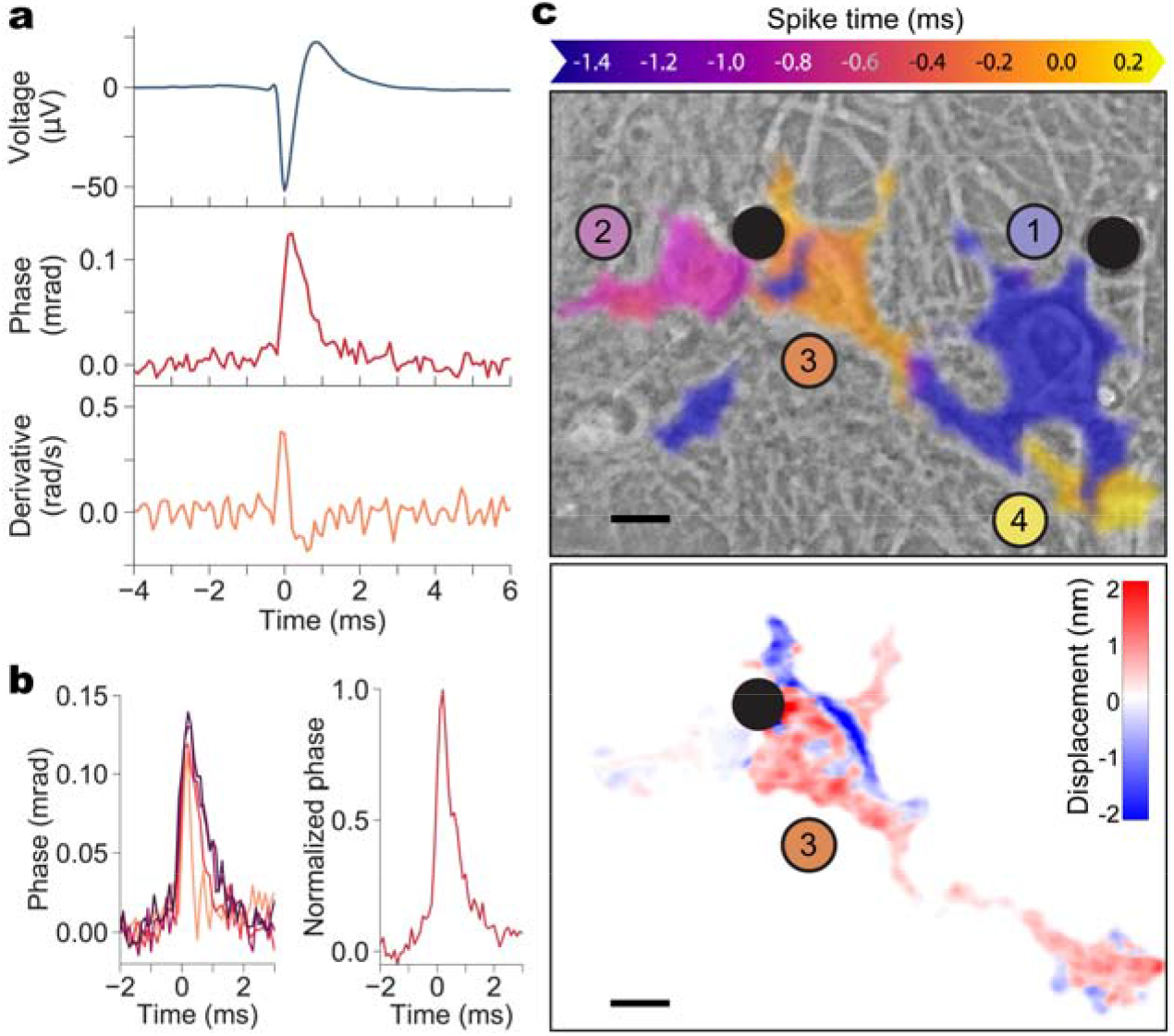
Synchronized QPI/MEA measurements of neuronal deformations. (a) Top: extracellular electrical recording used to synchronize video frames. Middle: spike-triggered average phase change across the cell during the action potential. Bottom: time derivative of the phase signal matches the electrical waveform with a negative sign. (b) Left: time course of the phase signals from 4 cells, showing a quick onset to 0.12 mrad (average) followed by a 1~2 ms relaxation. Right: averaging all the datasets yields a spiking template for matched filter detection of spikes. (c) The spiking template helps detect the spike timing across the field of view (top), exploiting temporal and spatial adjacency of spiking signals to segment individual cells from the background. The bottom panel shows the displacements on the third neuron at *t*=0 ms (peak of the spike). Scale bar: 10 μm.

Phase changes across 4 neurons are shown in Fig. 2b (left), all consistent with the expected temporal dynamics of the action potentials. Because the phase signals were similar across all measured cells, a characteristic template for the spike detection can be obtained by averaging across the datasets (Fig. 2b right). The spiking template was generated from only the areas of higher SNR in multiple cells. Using this template as a match filter on every pixel in the FOV across time improved the spike detection and helped reject noisy areas in the image. Precise detection of the spike time allows separation of adjacent cells that fired in succession a few tenths of a millisecond apart. The resulting segmentation, shown in Fig. 2c (top) and in Supplementary Movie 2, illustrates these four cells in a FOV spiking in sequence over the course of 2 ms. Each neuron was segmented, overlaid on a phase contrast view, and color coded to represent the spike time, demonstrating the signal flow as the action potential moves from one neuron, along its processes, to another neuron spiking later. The frame from the STA phase movie corresponding to *t*=0, the spike time of the third cell, is shown at the bottom of Fig. 2c.

Moreover, template matching allows computational interpolation of the temporal resolution of the spike imaging from our native 0.1 ms, defined by the camera, down to 20 μs, as previously demonstrated with genetically encoded voltage indicators^30^. As shown in Supplementary Movie 3, with this method, the fast action potential propagation can be predicted with a temporal resolution of 20 μs.

### Spatial features of the spike-induced deformations

The measurement throughput on the MEA is defined by the prevalence of neurons adjacent to electrodes and firing individually. Good optical isolation of a cell required for QPI imaging contradicts the requirement of a high cell density necessary for high spontaneous spiking rate. These contradicting requirements resulted in relatively low yield of the QPI recordings of spontaneous firing in neuronal cultures. Using optogenetic stimulation^22^ and interferometric recordings synchronized to the optical stimuli, we obviated the need for the MEA. This approach relied on stable stimulation at 5 Hz rate (>99% success in eliciting spikes) with a small jitter (~0.1 ms) between the elicited spikes and the optical stimulus, as characterized by earlier MEA measurements (N=3). With this synchronization, we did not have to record electrical waveforms, and therefore neurons could be cultured on coverslips instead of thicker MEAs, which enabled high-resolution confocal microscopy for 3D morphology mapping.

However, optical stimulation resulted in a thermal phase artifact due to absorption of green light by the phenol red in the cell culture medium. We measured the heating artifact in the background of each STA via spatial averaging, which was then subtracted from the STA phase movie (see Methods), resulting in undisturbed spike-induced phase changes, as shown in Fig. 3a and Supplementary Fig. S2. The thermal artifact matched the curve predicted by the finite element modeling (COMSOL) based on the temperature dependence of the refractive index of water^31^.

**Fig. 3.**
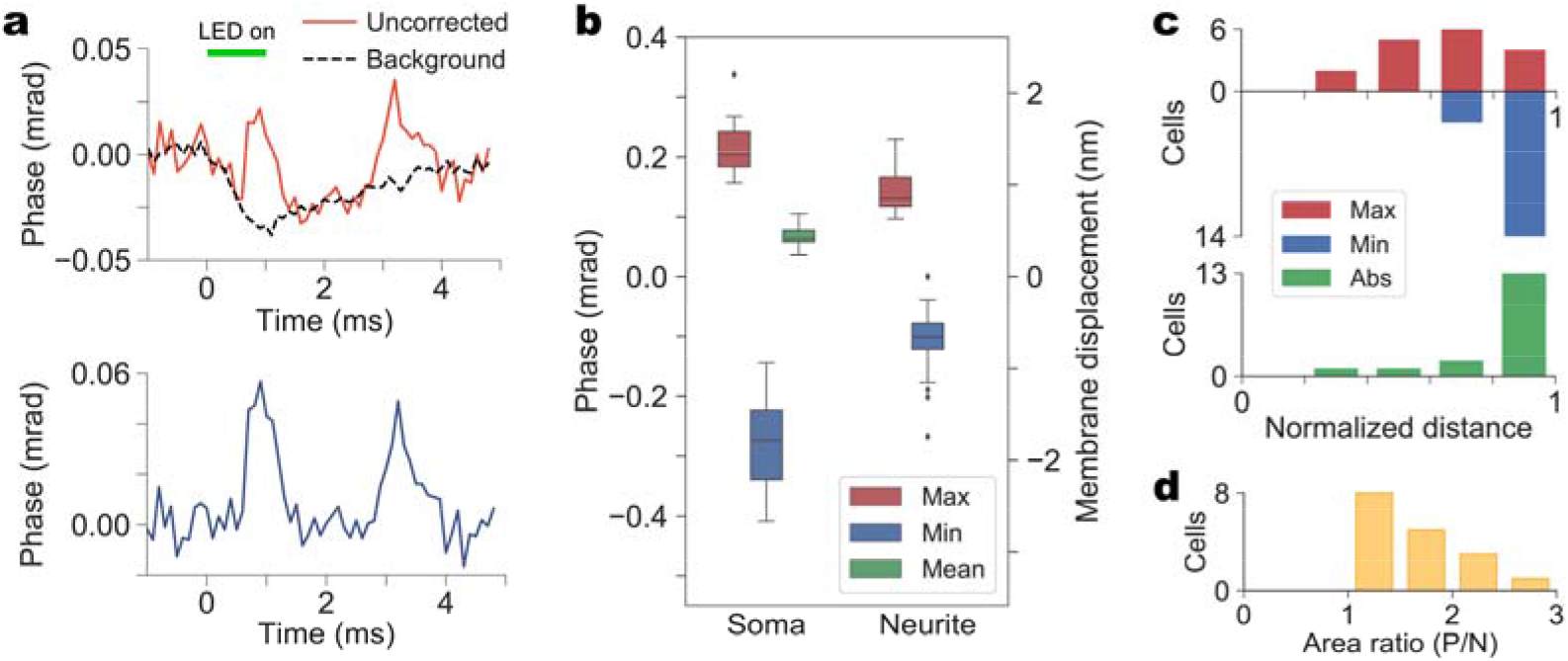
(a) Top: Thermal artifact from optogenetic stimulation overlaps with the spiking signal. Bottom: The same deformations after subtracting the thermal artifact. (b) Statistics of the displacements across cell somas (N=19) and neurites (N=34): maximum positive, minimum negative and mean absolute displacements. (c) Histograms of the lateral distances from the cell centroid to the locations exhibiting maximum positive displacements, minimum negative displacements and maximum absolute displacements (N=17). The distances are linearly normalized between the centroids (0) and the cell boundaries (1). (d) Histograms of the area ratios of the positive displacement regions to the negative displacement regions. No cells had a ratio below 1, meaning that areas of positive displacements are larger than the negative areas in every cell (N=17).

With the thermal artifacts removed, we acquired a large number of datasets imaging spike-induced deformations (N=19 for somas, N=34 for neurites). Figure 3b shows the magnitude of the deformations in cell somas and the surrounding neurites, quantified as a maximum positive displacement (1.3 ± 0.3 nm for somas, median ± standard deviation; 0.9 ± 0.2 nm for neurites), minimum negative displacement (−1.8 ± 0.6 nm for somas; −0.7 ± 0.3 nm for neurites) and the mean absolute displacement (0.4 ± 0.1 nm for somas). The latter was obtained by spatially averaging the absolute phase change over the entire soma.

In terms of spatial distribution, we observed a general trend of cells becoming more spherical during depolarization. An example of this is shown in Fig. 4. The top of the steep cell soma was moving down, while the periphery was getting thicker. A flat part of a neurite (indicated by an arrow) was also becoming more spherical: its middle portion was moving up while the edges were shifting down. Additional examples are shown in Supplementary Fig. S3. Intriguingly, we found that a large fraction (76%, N=17) of the maximum absolute displacements occurred in the regions near the cell boundaries, mainly contributed from the negative displacements (Fig. 3c); while every cell exhibited a larger positive displacement area than the negative displacement area (Fig. 3d, N=17). Theoretically, these observations are consistent with a general picture of cell centers moving up while boundaries move down, or cells shifting from one side to the other to become more spherical. From an engineering perspective, it also suggests that the boundary regions of neurons should be given a high priority for detecting the maximum signal for label-free imaging of action potentials.

**Fig. 4.**
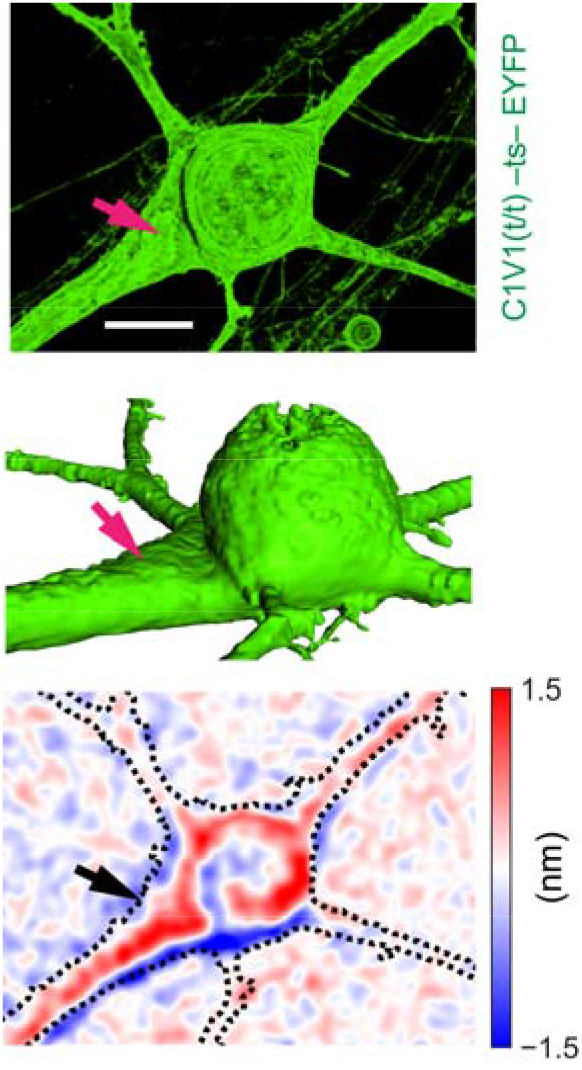
Confocal image (top) and its 3D perspective (middle), compared to the spike-induced deformation of a neuron (bottom). The arrow is pointing to a flat portion of the neurite which becomes more spherical, with its center moving up and the two boundaries shifting down. Scale bar: 10 μm.

### Modeling the voltage-induced cellular deformation

Our observations that the cells are becoming more spherical during membrane depolarization is in line with the voltage-dependent membrane tension model^15^, which predicts a quasi-linear relationship between the membrane tension and membrane potential:

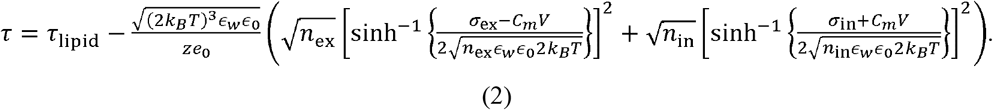

According to this model, the membrane tension should increase by ~5 μN/m for a 100 mV depolarization, given the surface charge density measured in neurons^32^.

Using the Young-Laplace law, the shape of biological cells is determined by the balance of hydrostatic pressure *p*_hydro_, membrane tension *τ*, and cortex tension *τ*_cortex_ ^23,33^. Membrane tension is composed of the lipid bilayer tension *τ*_lipid_ and the lateral repulsion of ions *F*_ions_ on both sides of the membrane. The mean surface curvature *H* on the cell surface can be determined as:

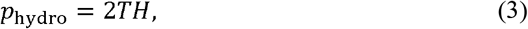

where

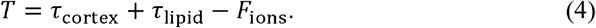

According to this model, increase in the surface tension Δ*T* during membrane depolarization will lead to an increase in the pressure towards the intracellular space, proportional to the surface curvature; i.e. areas with lower curvature will generate less pressure. For cell somas, this effect will drive the mean curvature on various parts of the cell to equalize, i.e., become more spherical.

To model the spike-induced deformation quantitatively, we applied the voltage-dependent membrane tension to a ‘cortical-shell liquid-core’ model of a single cell (Fig. 5a). The thickness and the Young’s modulus for the actin cortex were assumed to be 100 nm and ~1 MPa^25,34^, respectively. The viscosity of the cytoplasm was assumed to be ~2.5 mPa·s^35^ and the maximum membrane tension change was set to 5 μN/m. Perfectly spherical cells do not deform at all in this model, and the greater the aspect ratio and the overall size, the greater the displacements towards more spherical shape. This simple model yielded the same magnitude and distribution of positive and negative displacements as in our observations in cells.

**Fig. 5.**
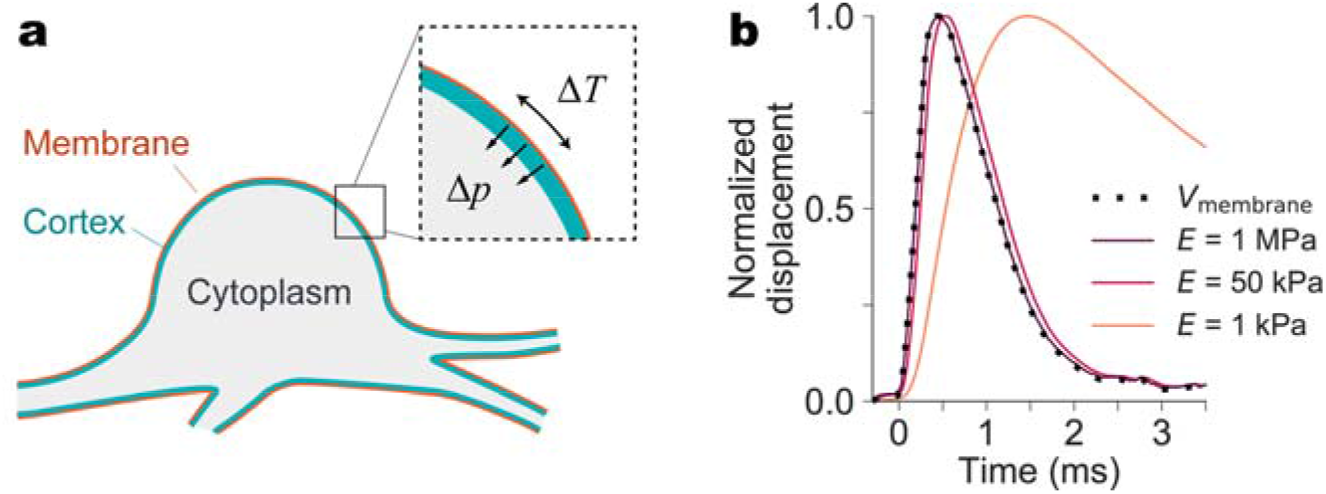
Mechanical model based on the voltage-dependent membrane tension predicts a deformation toward a more spherical shape during the action potential, with less than 0.1 ms latency. (a) The ‘cortical-shell liquid-core’ mechanical model of the cell transduces surface tension changes into cell deformation. (b) With a physiological Young’s modulus (1 MPa), predicted time course (~10 μs latency) matches the lack of observable delay between the deformation and electrical waveform. Much softer structures, however, would introduce significant delay.

This model links the response time, i.e. how quickly the spike-induced deformation will follow the action potential, to the cell stiffness (Young’s modulus). Figure 5b shows that with a Young’s modulus in the typical range from the literature^25,34,36^ (1 MPa), a cell responds to the tension change with a response time of ~10 μs (see Supplementary Fig. S4), tracking the underlying membrane potential closely, as we observe in experiments. In contrast, with low Young’s modulus (1kPa), the model predicts a delayed response, as the much softer membrane deforms the cell much slower, similar to earlier observations in *Chara* internodes with detached cell walls^37^. A Young’s modulus of 50 kPa shifts the deformation response past the 0.1 ms temporal resolution of our measurements, indicating that moduli below that limit would introduce a detectable delay relative to electrical pulse, which was not observed in our experiments.

Since the surface tension increase will make the cell more spherical, the deformation pattern should be largely dependent on the cell shape. To compare model predictions with experimental images, we acquired QPI and 3D confocal images of the same cell (Fig. 6a), and reconstructed its geometry for finite element modeling (Fig. 6b). Neurites were truncated as indicated by the white outline in Fig. 6a, and as shown in a 3D view in Fig. 6b. For a QPI imaging model, we incorporated the halo effect, an artifact due to the incomplete low-pass filtering of the reference beam^38,39^ (See Supplementary Material). The measured and predicted deformation patterns, shown in Fig. 6c, d, exhibit similar magnitudes of displacements along with a general match of features, such as the negative region at the top of the soma and the band of positive displacements in a crescent through the middle. Additional examples are shown in Supplementary Fig. S5 and Supplementary Movie 4.

**Fig. 6.**
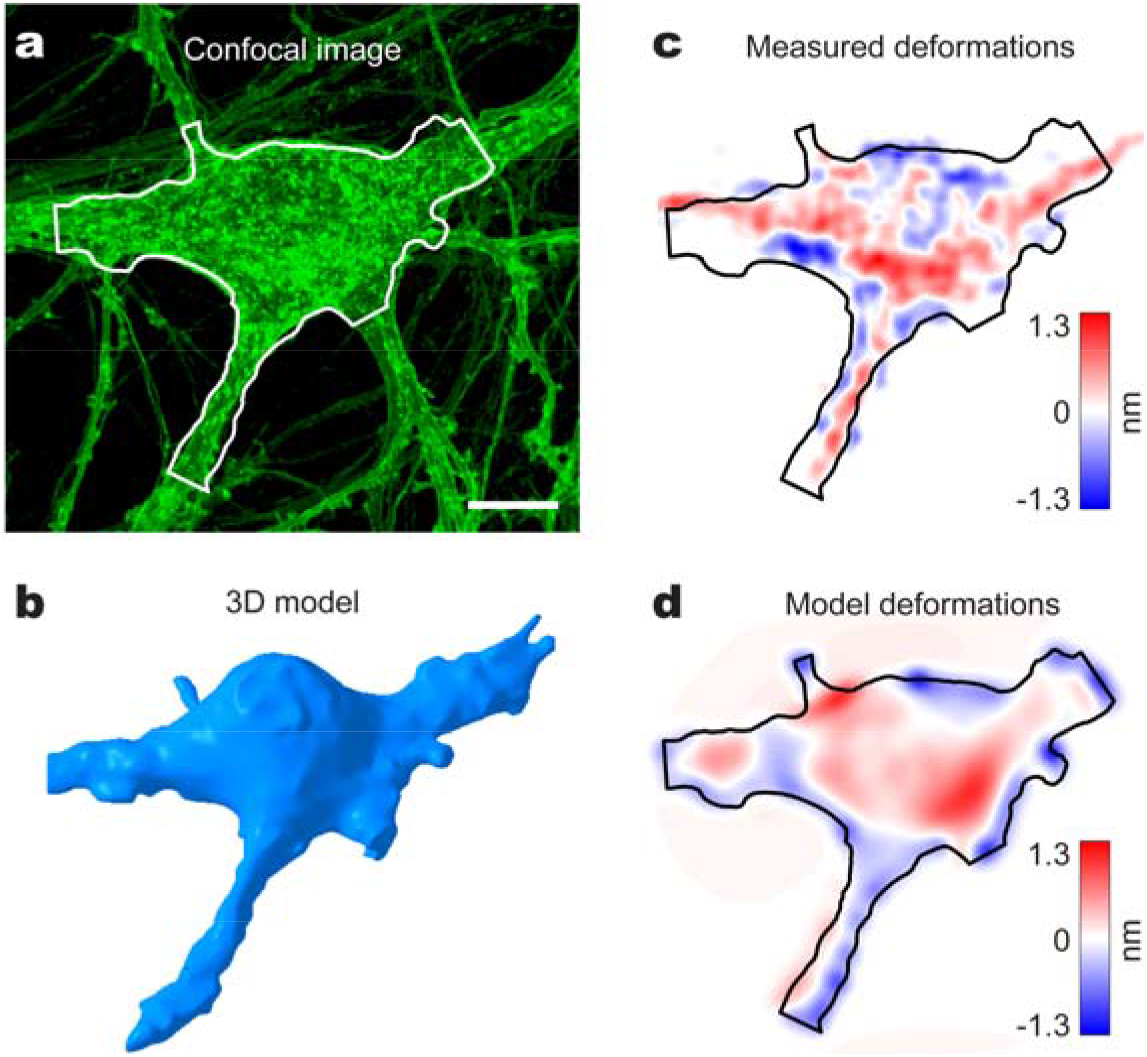
Comparison of the modeled and experimental spike-induced deformations in the same cell. (a) Confocal image of a cultured neuron with the segmented area outlined in white. Scalebar: 10 μm. (b) 3D model of the cell segmented from the confocal image stacks. (c) QPI of the spike-induced deformation of the same cell. (d) Simulated deformations from the model, combined with a QPI imaging model that incorporates the halo effect and the refractive index difference.

## Discussion

Interferometric detection of the spike-induced deformation offers a fast, intrinsic probe for label-free imaging of action potentials. Exogenous fluorescent indicators require invasive interventions to load the cells with optical markers and suffer from photobleaching^40^ and heating limitations^30^. In addition, the slow response of calcium imaging (>100 ms decay^41^), makes it inadequate for revealing closely spaced action potentials. Response time of the spike-induced deformation reported here (<0.1 ms) is much better than even the genetically encoded voltage indicators, the response times of which are on the order of 1 ms^41,42^.

However, interferometric imaging of single action potentials in neurons calls for further SNR enhancement. This could be addressed using reflection geometry (like optical coherence tomography, OCT), in which the signal benefits from the double pass and is not attenuated by the refractive index difference between the cell and the medium, as in transmission QPI. Given a refractive index difference around 0.02 and a refractive index of 1.33 for water, a 133-fold improvement is expected in the phase change during the spike (about 13.3 mrad) in reflection mode. In OCT, this phase change corresponds to the phase noise floor at a reasonable SNR of ~38 dB^43^. The key question, though, is the strength of the backscattered signal. At the medium-cell boundary, the reflection coefficient is proportional to the square of the refractive index difference. Therefore, the SNR, which scales reciprocally to the square root of the number of photons, will be attenuated by the refractive index difference, exactly negating the benefit of stronger phase change in reflection for a given laser power. However, if the scattering coefficient is enhanced by the turbulent tissue properties or by contrast enhancement agents^44,45^, the SNR might be significantly increased.

Another interesting possibility is to improve SNR using template-matching techniques based on full spatial and temporal correlations in the 4D spike-induced deformation movie. These could include machine learning algorithms, along with spatial averaging over the ROI predicted by the 3D cell model from confocal microscopy or holographic tomography^9,46^.

Full-field imaging of cellular deformations at high temporal and spatial resolution enabled comparison of the experimental data with 3D modeling to elucidate the underlying mechanism of electromechanical coupling in cells. This comparison demonstrated that the voltage-dependent membrane tension-based model correctly describes the magnitude of the effect, the time course and its spatial features. The main distinction of our model from the report from El Hady *et al.*^10^ is that we assume the voltage-to-force relation to be quasi-linear rather than squared. This is because in previous experiments with HEK cells, no second harmonic has been observed in the frequency response^18,19^, as would be expected if the relationship would be quadratic.

Given the non-spherical shape of neurons in the brain^47,48^, we expect deformations to be present, such that non-invasive label-free imaging of action potentials, especially in reflection geometry, may offer invaluable opportunities for real-time, high-bandwidth read-out from large areas in the brain. Our observations of the spike-induced cellular deformations and validation of the underlying theory bring this goal closer to practical implementation.

## Acknowledgements

Funding was provided by the NIH grant U01 EY025501 and by the Stanford Neurosciences Institute.

We would like to thank Drs. A. Roorda, B.H. Park, R. Sabesan, E.J. Chichilnisky for fruitful discussions, and R. Tien, C. Raja for helping with dissection of cortical tissues for neuron culture.

## Author contributions

T.L., K.B. and D.P. designed research. T.L. and K.B. performed experiments. T.L. and K.B. performed computational modeling. C.R. and K.D. provided cortical tissue and insights on opsin selection. V.Z. and T.L. prepared cell cultures. T.F. conducted the confocal imaging. T.L. and K.B. analyzed the data. T.L., K.B. and D.P. wrote the manuscript. All work was supervised by D.P.

## Methods

### Primary neuronal culture

All experimental procedures involving animals were conducted under protocols approved by Stanford University Administrative Panel on Laboratory Animal Care. Primary neuronal cultures were isolated from postnatal day 0 (P0) rat cortical tissues. The cortices were dissected, the meninges removed, and tissue was digested with 0.4mg/mL papain (Worthington Biochemical Corporation) in Hanks’ balanced salt solution (HBSS, Cat #14175-095, Gibco) at 37 °C for 20 minutes. The digested tissue was then triturated, and cell pellet was resuspended in Neurobasal-A medium (Gibco), supplemented with 2mM GlutaMax (Gibco), NeuroCult SM1 Neuronal Supplement (1:50 dilution, Stemcell Technologies) and 5% fetal bovine serum (Sigma-Aldrich). Cells were plated on multi-electrode arrays (MEAs) or the gridded imaging dishes with coverslip bottoms (μ-Dish 35 mm, high Grid-500, ibidi), pre-coated with poly-D-lysine (PDL) (0.1mg/mL, Sigma-Aldrich). The next day the plating medium was replaced with serum–free neuronal culture medium (Neurobasal-A (Gibco) medium, 2mM GlutaMax (Gibco), NeuroCult SM1 Neuronal Supplement (1:50 dilution, Stemcell Technologies). Primary neuronal cultures were kept in humidified incubator with 5% CO_2_ at 37 °C and maintained by half volume culture media exchange every 3-4 days.

For optogenetic stimulation experiments cells were plated at a density of 500-750 cells/mm^2^, transduced with pAAV-CaMKIIa-C1V1 (t/t)-TS-EYFP^1^ (AAV9) at 5-7 days in vitro (DIV5-7) (Addgene viral prep # 35499-AAV9; http://n2t.net/addgene:35499; RRID:Addgene_35499; Plasmid was a gift from Karl Deisseroth) and supplemented with 10μM fluorodeoxyuridine (FUDR, Sigma-Aldrich) solution to control glial cell growth^2,3^. For spontaneously spiking cultures, the cells were plated at density of 1,200-2,000 cells/mm^2^ on MEAs, pre-coated with Matrigel, in addition to poly-D-lysine (PDL). Quantitative phase imaging (QPI), electrical recordings (for MEAs) and live-cell confocal microscopy were performed on DIV17-28.

### Recording of the spike-induced deformations

#### Quantitative phase microscopy

Quantitative phase microscopy based on common-path interferometer^4,5^ was adapted from our previous work^6,7^ (Supplementary Fig. S1). Light from the superluminescent diode (SLD, SLD830S-A20; Thorlabs) was collimated by a fiber-coupled collimator (F220FC-780; Thorlabs). Neurons on MEAs or on imaging dishes were imaged via 10X objective (CFI Plan Fluor 10X, N.A. 0.3, WD 16.0 mm; Nikon), and the light beam recollimated by a 200 mm tube lens (Nikon) was projected through a transmission grating (46-074, 110 grooves/mm; Edmund Optics) and a subsequent 4-f system. The 0^th^ diffraction order of the grating was filtered through a 150 μm pinhole mask on the Fourier plane, while the 1^st^ order was passed through without filtering. The 4-f system consisted of a 50 mm lens (AF Nikkor 50mm f/1.8D; Nikon) and a 250 mm bi-convex lens (LB1889-B; Thorlabs). The camera sensor (Phantom v641; Vision Research) captured the resulting interferograms at a frame rate of 10 kHz and a resolution of up to 768×480 pixels. For optimal data transfer, resolution was reduced to 512×384 pixels in most datasets.

#### Extracellular electrical recording using MEA

Action potentials were recorded extracellularly using a custom 61-channel MEA system built on a transparent substrate with ITO leads^8^. The recording electrodes were electro-deposited with platinum black prior to culturing the neurons on MEAs. The cell growth areas on MEAs were 2 cm^2^ in size. Recorded electrical signals were amplified with a gain of 840 and filtered with a 43 - 2000 Hz bandpass filter. A data acquisition card (NI PCI-6110; National Instruments) sampled the signals at 20 kHz. To synchronize the QPI recording with the MEA system, an external clock signal from a function generator connected to the camera’s F-Sync input was repeatedly triggered from the MEA system every 100 ms. In addition, the ready signal from the camera, marking the start of an image sequence acquisition, was recorded by the MEA system to synchronize the electrical and optical recordings.

#### Optogenetic stimulation

A fiber-coupled LED (M565F3; Thorlabs) with its center wavelength at 565 nm was used to deliver light stimulation to the neurons expressing C1V1 (t/t) opsins. Output of that fiber was coupled to a collimator (F260SMA-A; Thorlabs) and the beam aligned into the rear port of the microscope (TE2000-U; Nikon) was directed onto the sample via dichroic mirror (FF585-Di01-25×36; Semrock) (Supplementary Fig. S1). Through the 10X objective, the beam was focused onto the sample plane to a ~300 μm spot with a power density of ~5.5 mW/mm^2^. Illuminated spot was coaligned with the FOV of the QPI, so that the neurons of interest could be stimulated optically. An optical bandpass filter (FF01-819/44-25; Semrock) was placed in the imaging path to further filter out the back-reflected green light from the QPI beam.

#### Selective recording of the spike-induced deformation

To record spike-induced deformations from spontaneously firing neurons, real-time spike detection was utilized to selectively search for and record the burst firings, while neglecting other sparse action potentials which wastes limited recording time and data storage. The camera was set to record 5000 pre-trigger frames and 1 post-trigger frame at a frame rate of 10 kHz. Once the MEA system detected more than a predetermined number of spikes (usually 10) in a 500-ms window, a trigger was sent automatically to the camera to save the previously buffered 500 ms of interferograms from the camera’s random-access memory (RAM) to its nonvolatile memory (CineMag II 512 GB; Vision Research). For optogenetically stimulated neurons, selective recording was implemented by capturing only the following 5~9-ms movie segments after the 5-Hz stimuli, along with the preceding 1-ms segments, and saved directly to the computer hard disk.

### Data processing

#### Spike-triggered averaged (STA) phase movie

The phase differences between the first interferogram in each movie sequence and the subsequent frames were computed by the fast phase reconstruction algorithm for diffraction phase microscopy^9^, with additional acceleration using a graphics processing unit (Tesla K40c; Nvidia). Global phase fluctuation in each frame was eliminated by subtracting the spatial average of the background, while highly noisy pixels were excluded from the background ROI.

The STA phase movie was generated by aligning the phase difference movies to their spike times and averaging them together. The slow lateral drifts between movies were corrected by monomodal image registration based on the QPI images of their first frames. To ensure optimal averaging efficiency, noisy phase movies were rejected to prevent unexpected events, such as moving particles or sudden vibrations, from affecting the STA results.

#### Heating correction

Heating correction was applied to the optogenetically stimulated datasets. The phase artifact due to heating from the LED stimulation has two unique features: 1) predictable time dynamics that matches the finite element modeling (FEM) based on the temperature dependence of the refractive index of water, 2) low spatial frequencies compared to the cell structure. Leveraging these features, the heating signal can be decoupled from the phase change of the spike-induced deformation. First, a 50×50-pixel spatial averaging filter was applied to the STA phase movie to generate a spatially smoothed phase movie. The heating ROI was extracted by thresholding the cross-correlation between the template predicted by the FEM and the phase change on each pixel of the smoothed phase movie (Supplementary Fig. S2). The measured heating curve was obtained by spatially averaging the detected heating ROI while excluding the spiking ROI mentioned below. This curve, unique to each dataset, was then used to fit the phase change due to the heating effect on each pixel in the smoothed phase movie by linear regression with normal equation, while neglecting the sample points when the cells spike. The heating phase change fit to each pixel was subtracted from the phase change on that pixel in the original STA phase movie (Supplementary Fig. S2).

#### Spiking ROI segmentation

The distinct spatial and temporal clumping of the phase signal in pixels throughout the STA was used to enhance the detection and segmentation of spiking cells compared to the shot-noise limited performance of single pixels. By referring to previous observations of the spiking signal and its relation to the mechanical deformation, we used a known template to match-filter each pixel. The output of this correlation gave an indication of how likely the spiking signal occurred on this pixel throughout the duration of the STA, as well as when the spike occurred, but was insufficient to segment every spiking pixel from the noisy background. We consolidated this per-pixel information across the whole STA to clump nearby pixels that spike together into a single spiking cell. We then grew those regions to fill holes left by the noise, according to the absolute phase measurement that indicated the cell structure. The result was a 3D spatiotemporal mask that passed only those pixels of the STA that had been detected as spiking at a given location and time (Supplementary Movie 2).

### Confocal image acquisition

3D imaging was performed using a Zeiss LSM 880 inverted confocal microscope with Zeiss ZEN black software (Carl Zeiss). Imaging dishes with cultured cells were placed on the microscope stage inside a large incubator (Zeiss Incubator XL S1) kept at 37 °C during imaging. Image planes were acquired through the total thickness of the cell using a Z-stack, with upper and lower bounds defined 3 μm below the cover slip and above the cell membrane. Stacks were acquired using a 63x/1.4 NA oil immersion objective (Plan-Apochromat 63x/1.4 Oil DIC M27; Carl Zeiss) with 135 μm × 135 μm acquisition area, 447 nm z-step, and 930 nm pinhole, with n=8 line averaging. Deconvolution on the acquired image stacks was performed using Zeiss Zen blue software (Carl Zeiss) using the fast iterative method. For illustrating 3D cell shapes in Fig. 4 and Supplementary Fig. S3, images were analyzed using Imaris software (Bitplane).

### Reconstructing 3D cell shapes from confocal images

The confocal image stacks were manually segmented in 3D Slicer^10^ (http://www.slicer.org) and exported as meshes to Netfabb software (Autodesk) to be converted into parametric surfaces for import into FEM software (COMSOL). Semi-automated segmentation algorithms, such as region growing and volume dilation, were combined with user-generated seeds and editing to select the cell of interest, including its nucleus and processes as far out as could be distinguished. Smoothing and remeshing operations were performed before generating a smooth parametric surface as the best fit to the mesh.

### Modeling the voltage-induced cellular deformation

FEM was implemented to model the cellular deformation due to the change in voltage-dependent membrane tension, using COMSOL Multiphysics software (v5.4). Based on the ‘cortical shell-liquid core’ model, we assumed a 100-nm thick cortical shell wrapping around a hemispheroidal liquid core. The cortical shell was set to be a linear elastic material since the cellular deformation is minute. A Young’s modulus of ~1MPa was calculated from the previous reports using the power-law relation due to viscoelasticity^11–13^. The liquid core was assumed to be an incompressible laminar flow with a non-slip boundary condition. The viscosity of the liquid core was set to 2.5 mPa·s. To simulate the spike-induced deformation, a voltage-dependent membrane tension of 5 μN/m was applied to the external surface of the cortical shell.

To compare the predicted deformations with the phase imaging results, 3D cell shapes were imported into COMSOL using CAD import module. Except for the cell geometries, other parameters were set to be the same as those in the hemispheroidal model. Modeled phase images of cells were extracted using general projection operation through the *z* axis, while additional halo effect in the QPI and spatial blurring due to the limited optical resolution were applied to the forward modeling.

