## Supplementary Information for "How neurons move during action potentials"

**Supplementary Figures**

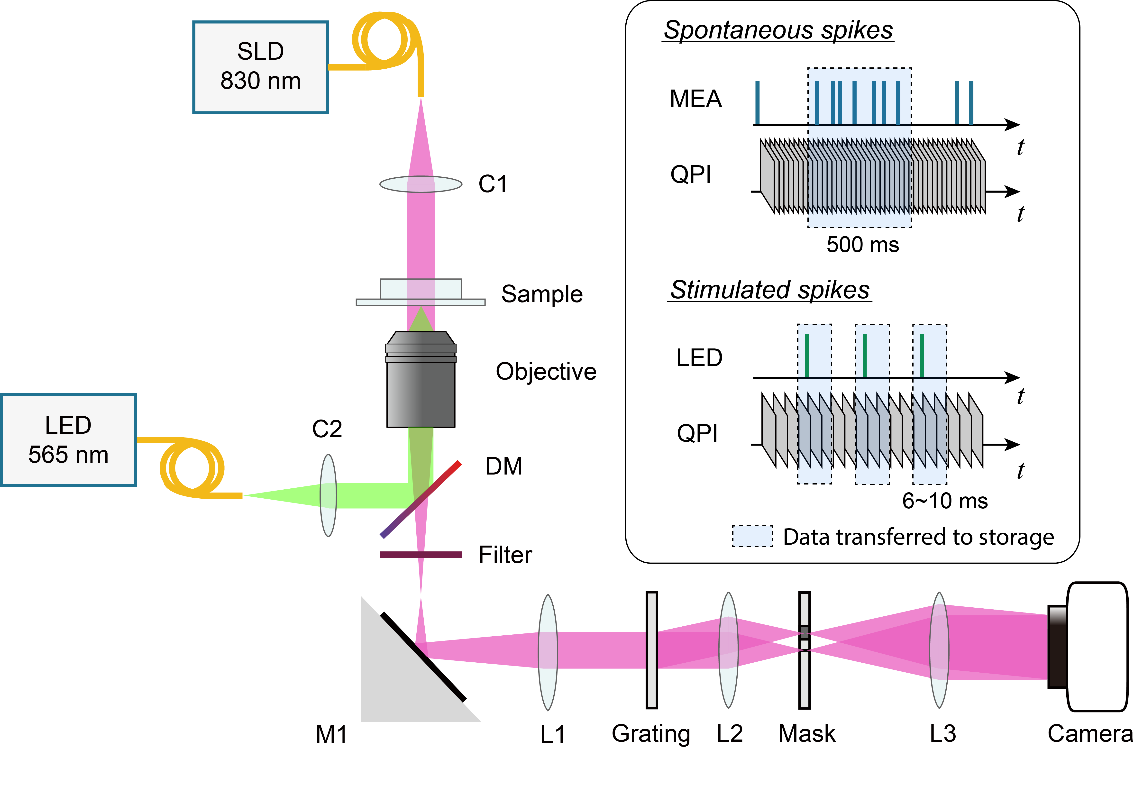

**Figure S1:** Experimental setup of the quantitative phase microscopy and recording strategies for spontaneously firing and optogenetically stimulated neurons (C1, C2: collimator; DM: dichroic mirror; M1: mirror; L1, L2, L3: lens). For spontaneously firing neurons, spikes are detected in real time and 500-ms movie segments containing the burst firing are saved to the nonvolatile memory. For optogenetically stimulated neurons, the 6~10-ms movie segments around the stimulated spikes are saved.

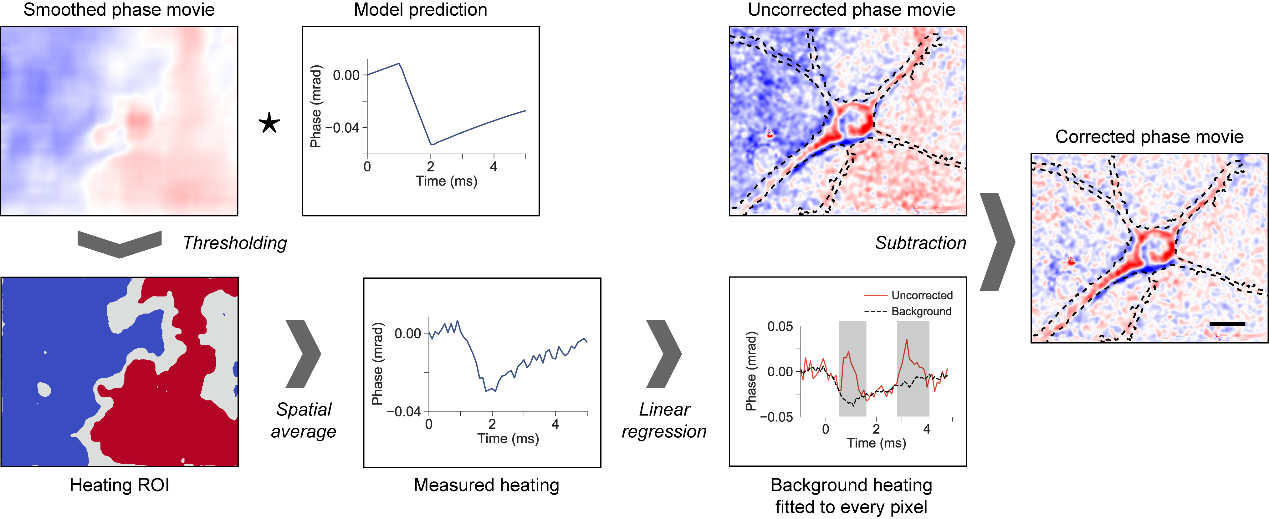

**Figure S2:** Flowchart of the heating artifact correction. A heating ROI is extracted by thresholding the cross-correlation between the smoothed phase movie and the modeled template, and the measured heating is obtained by spatially averaging the heating ROI while excluding the ROI of the spiking cells. This curve is used to fit to the phase changes in time, except for that during cell spiking (marked by gray regions), on each pixel of the smoothed phase movie using linear regression. Subtracting the fit heating artifacts from the original STA phase movie results in a corrected phase movie.

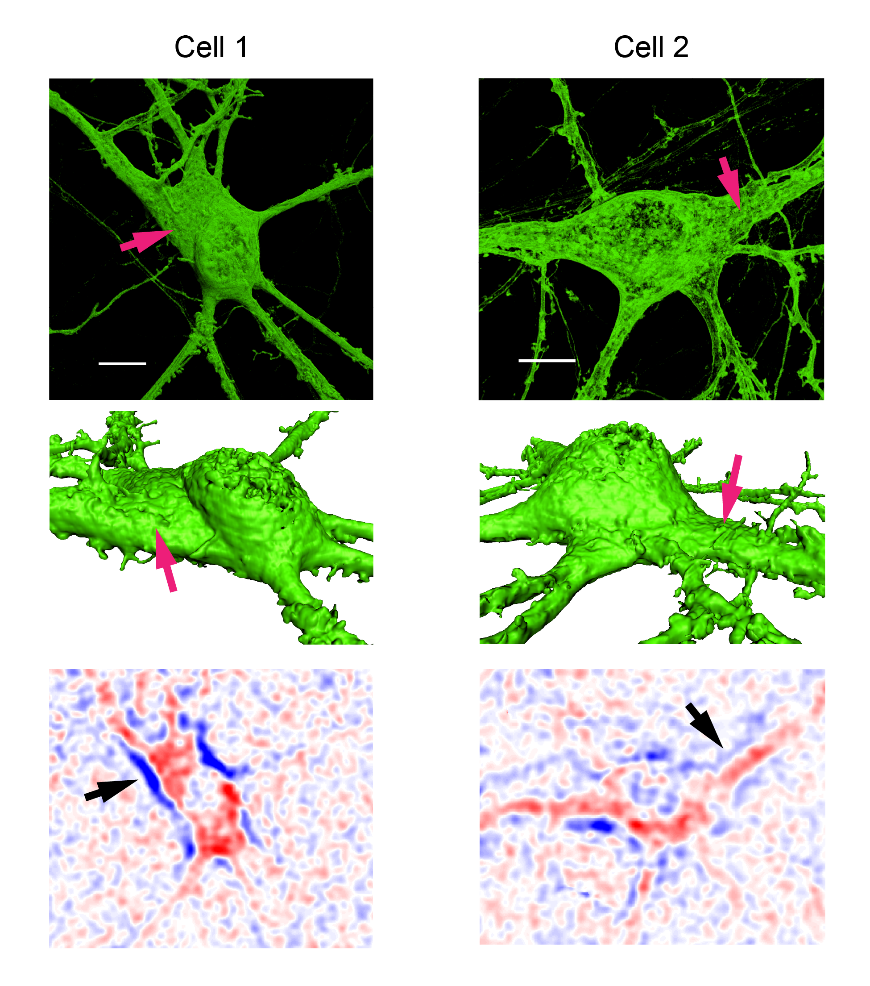

**Figure S3:** Additional examples illustrating how the flat part of the cell becomes more spherical. Arrows indicate a flat part in each cell, which is moving up in its center and down along the boundaries. Scale bar: 10 μm.

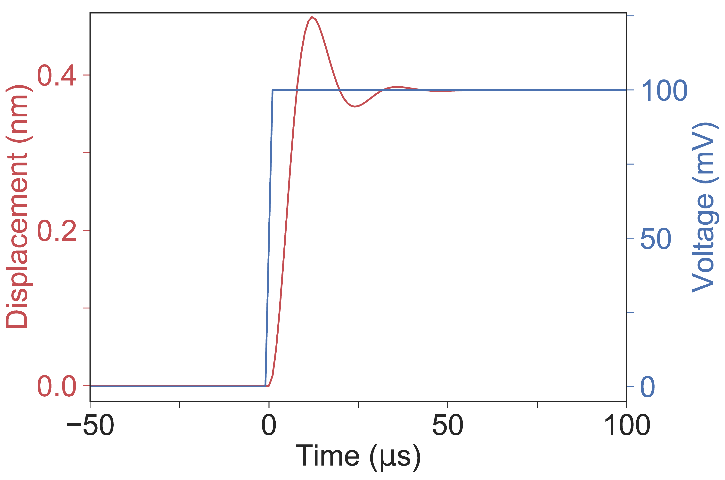

**Figure S4:** Modeled time course of the hemispheroidal cell deformation in response to a 100 mV voltage step. Response time is approximately 10 μs.

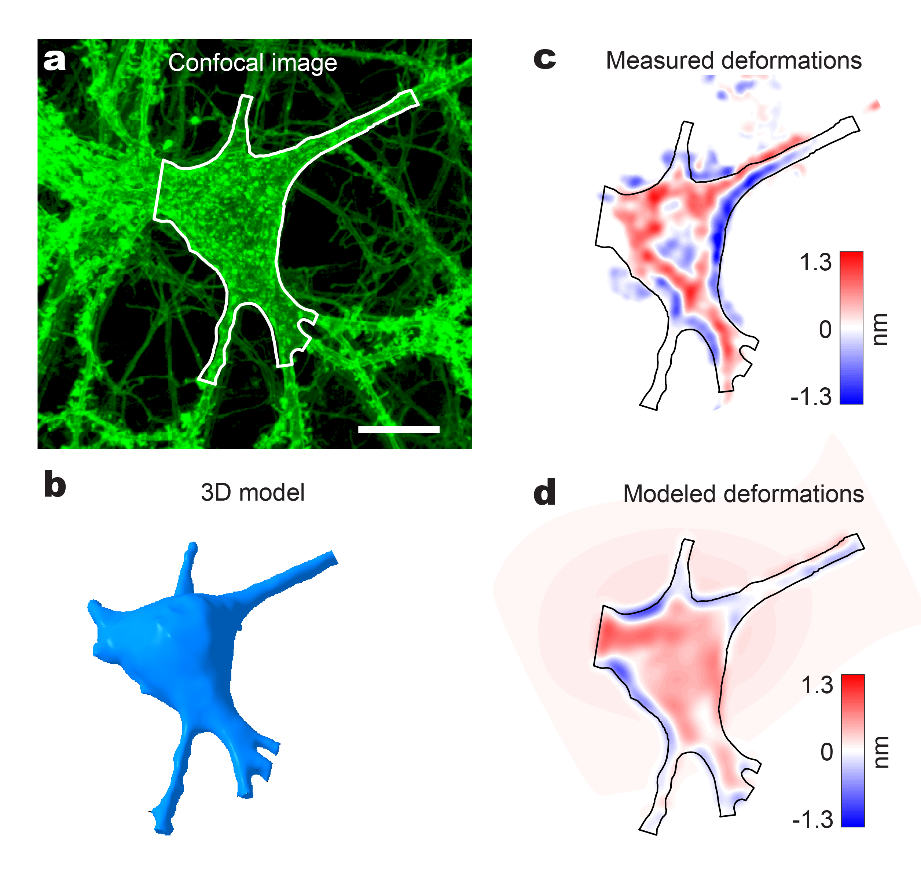

**Figure S5:** Comparison of the modeled and experimental spike-induced deformations in an additional cell. (a) Confocal image of a cultured neuron with the segmented area outlined in white. Scale bar: 10 μm. (b) 3D model of the cell segmented from the confocal image stacks. (c) QPI measured spike-induced deformation from the same cell. (d) Simulated deformations from the model, combined with a QPI imaging model that incorporates the halo effect and fits the refractive index difference.

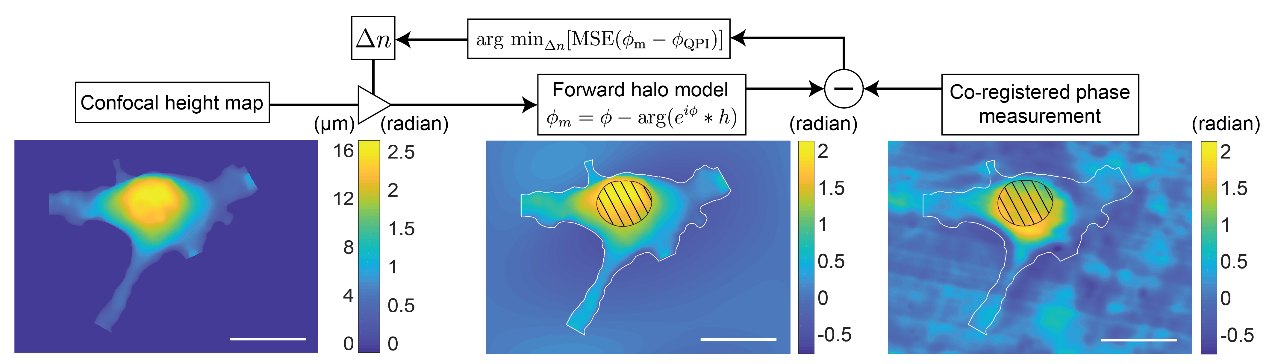

**Figure S6:** Block diagram of the refractive index calibration procedure. (Left) the confocal height map is converted into radians for a range of possible $\Delta n$. (Center) each phase image is then passed through a forward halo model based on the size of the pinhole in the imaging system. (Right) the measured phase image is compared to the forward modelled phase image derived from the confocal measurement, and the mean squared error (MSE) over the soma region is used to find the best $\Delta n$ fit. The nucleus is segmented from the confocal measurement, as indicated by black bars, and is excluded from the fit region, as are the areas along the white boundaries of the confocal image, which contain artifacts from the forward modelling due to abrupt transitions.

**Supplementary Movies**

**Movie 1:** Spike-triggered average (STA) phase movie of spontaneous action potentials in a cell on the MEA with a FOV of $106 \mu m\times80 \mu m$. Scale bar: 20 μm.

**Movie 2:** Original and processed views of 4 spiking cells. (Left) the SNR movie shows the signal to noise ratio in each pixel, with the cell deformations appearing in the presence of background noise. (Right) the processed phase movie shows the same four cells after they have been segmented from the background by the spiking ROI segmentation (see Methods). Scale bar: 20 μm.

**Movie 3:** Super-resolution timing prediction showing the 20 μs time resolution propagation of a spike across a cell. The native 100 μs time resolution is interpolated in order to more precisely determine spike time across local regions of the cell by detecting the peak correlation with the spiking template at a higher sampling rate than actually measured. Scale bar: 20 μm.

**Movie 4:** Visualization of the cellular deformations induced by the changing cell potential. Middle frame demonstrates the computed changes in the optical pathlength through the cell. The right frame shows 3D rendering of the cellular deformations amplified 2000x for better visibility of the minute cellular movements.

**Supplementary Information**

***Refractive index calibration***

The optical phase change of the light passing through the sample relative to the background is related to the height $h$ of the sample as following:

$$h=(\lambda\cdot\Delta\varphi)/(2\pi\cdot\Delta n)$$

where $\Delta\varphi$ is the phase change measured at wavelength $\lambda$, and $\Delta n$ is the difference of refractive index between the sample and the surrounding medium. In principle, calibration of the $\Delta n$ could be done if the height of the object and the phase difference are known from two independent measurements, such as QPI and confocal imaging. In the case of QPI with a finite-size pinhole, the halo-effect adds an image-dependent artifact to structures of sufficiently low spatial frequency. The halo effect can be computed as,

$$\phi_{m}=\phi-\arg\left( e^{i\phi}*h \right),$$

where $\phi$ is the underlying phase distribution, $h$ is the spatial domain representation of the pinhole, in our case a 150 μm diameter filter, $*$ is the convolution operator, and $\phi_{m}$ is the finaly observed phase distribution with a halo artifact. A QPI system with an infinitely small pinhole (a delta function at the Fourier plane of the QPI system) would measure the underlying phase perfectly, but this would transmit no light and hence no interferogram could be formed. In practice, a finite pinhole must be sized to balance the amount of light in the reference and sample arms of the common-path QPI system. Therefore, image information below a certain spatial frequency will be transmitted through the pinhole and will generate artifacts in the interferogram. This results in a halo around the phase objects^1,2^.

To estimate the halo effect and extract the correct $\Delta n$ by comparison with the experimental image, we pass the confocal height map through a forward imaging model sweeping across $\Delta n$ (Supplementary Fig. S6). For each $\Delta n$, the result is compared to the measured phase, and the mean squared error over the soma region is used to find the best $\Delta n$ match. Since the refractive index of the cell nucleus is another unknown, we segment the nucleus out. Using this procedure, an estimate of $\Delta n=0.02$ was obtained in two cells, and taken as the refractive index difference for calculating the height from the phase in all other datasets.

1	Nguyen, T. H. *et al.* Halo-free Phase Contrast Microscopy. *Scientific Reports* **7**, 44034 (2017).

2	Kandel, M. E., Fanous, M., Best-Popescu, C. & Popescu, G. Real-time halo correction in phase contrast imaging. *Biomed. Opt. Express* **9**, 623-635 (2018).
